# Timing Determines Fate: Temporal Dynamics of Jasmonate Signaling Distinguish Adaptive from Maladaptive Salt Stress Responses in Rice

**DOI:** 10.64898/2026.10.02.756191

**Authors:** Nitin Pawar, Michael Riemann, Elisabeth Eiche, Peter Nick

**Affiliations:** Molecular Cell Biology, Joseph Kölreuter Institute for Plant Sciences (JKIP), Karlsruhe Institute of Technology (KIT), Fritz-Haber-Weg 4, 76131 Karlsruhe, Germany; Department of Crop Sciences, Viticulture & Oenology Group, Research Institute of Organic Agriculture (FiBL Switzerland), Ackerstrasse 113, CH-5070 Frick, Switzerland; Institute of Applied Geosciences (AGW), Karlsruhe Institute of Technology (KIT), Adenauerring 20b, 76131 Karlsruhe, Germany

**Keywords:** Allene oxide cyclase, jasmonate signaling, potassium transport, rice (*Oryza sativa*), salinity tolerance, salt stress

## Abstract

Plant encounters with saline conditions trigger a phase of osmotic stress, followed by ionic stress as salt accumulates in plant tissue. Jasmonate (JA) signalling plays a vital role in both phases, but its role for adaptation depends on timing. Using wild-type rice (*Oryza sativa* L. cv. Kitaake) and a JA-deficient allene oxide cyclase (*aoc*) mutant, we followed whole-plant physiology (growth, biomass, photosynthesis, ion distribution) with expression analysis for genes of jasmonate signalling, ion transporters and redox balance to separate adaptive from maladaptive JA effects. The *aoc* mutant sustained shoot and root growth more effectively than the wild type, maintaining cell expansion in the shoot and cell proliferation in the root under saline conditions. Likewise, stomatal conductance in the mutant was more persistent under salinity, accompanied by improved shoot K /Na homeostasis and reduced sodium in young leaves. In the wild-type, prolonged saline conditions were linked with induction of *OsJAZ13* and *OsNHX1*, stomatal closure and chlorophyll degradation. We discuss these findings in the context of a model, where the function of JA signalling does not merely depend on its accumulation, but on its temporal dynamics. Here, *OsHAK16* emerges as a marker of early adaptive responses, whereas *OsNHX1* and *OsJAZ13* characterize later maladaptive stress states, emphasising the potential of modulating JA dynamics to improve salt resilience in rice breeding.

**Highlight:** Time-resolved physiology and transcript profiling show that jasmonate signalling promotes salt tolerance in rice when transient but drives ionic damage when sustained.

## Introduction

Increasing salt concentrations have become a major threat to global food production. The proportion of irrigated agricultural land affected by salinity is currently around 20%, and it is increasing day by day due to the use of brackish water and excessive evaporation in consequence of non-sustainable irrigation. The challenge is accentuated by global climate change, leading to sea-level rise, but also unpredictable rainfalls (Tarolli *et al*., 2024). Salinity stress comes in two waves. As an immediate effect of exposure to salt, the plant suffers from turgor loss, such that growth is stalled. This osmotic stress is followed by ionic stress, when sodium ions enter the cell through Non-Selective Cation Channels and perturb ion homeostasis, electron transport in mitochondria and chloroplasts, as well as the activity of important metabolic enzymes (for review see Munns and Tester, 2008). To breed for new crops with better resilience to salinity requires a thorough knowledge of the mechanisms underlying stress damage but also stress adaptation.

Rice belongs to the glycophytes, i.e., plants that are susceptible to salinity. Already 40-50 mM NaCl can cause substantial reductions in growth and grain yield. This salt susceptibility seems to be, at least partially, linked with drawbacks in ion transport. Unlike other crops, such as sorghum (Abuslima *et al*., 2022), rice is unable to retain sodium in the roots, such that Na^+^ ions can easily pass the endodermal barrier and are then translocated to the shoot via the xylem in an unrestrained manner (Yeo *et al*., 1987), causing a strong depletion of potassium from the shoot. A central component of plant salinity tolerance is the regulation of ion transport across cellular membranes. For rice, salt resistance can be at least partially attributed to the activity of such transporters. For instance, a more efficient variant of the high affinity K^+^ transporter (HKT) family that can load shoot sodium into the phloem and unload it in the root, was shown to mediate the resistance of the salt-tolerant *indica* landrace Nona Bokra (Ren *et al*., 2005). Later, this transporter was identified as OsHKT1-5 (Kobayashi *et al*., 2017). In addition to HKT transporters, the sodium exporter Salt Overly Sensitive 1 (SOS1) has been found to be crucial. This transporter is activated by the calcium- sensitive complex of its regulators SOS2 and SOS3 that can sense the rapid influx of calcium triggered by the osmotic component of salinity (reviewed in Ji *et al*., 2013). A loss-of- function mutant of *SOS1* not only accumulates more sodium in the root, but also partitions more sodium to the shoot, indicating that this transporter is also involved in removing sodium from the xylem (El Mahi *et al*., 2019). Interestingly, this mutant exhibited only a dampened response of salinity-induced genes in the root, although the sodium content was elevated, indicating that SOS1, in addition to act as sodium excluder, has a function in sensing salinity. A third group of transporters acts at the vacuole. These sodium-proton exchangers (NHX) can sequester sodium in the vacuole, thus, reducing its cytosolic concentration, while preserving turgescence and, thus, the ability to grow. In fact, overexpression of *OsNHX1* was shown to improve salt tolerance in rice (Fukuda *et al*., 2004).

The activity of these transporters must be regulated immediately, meaning that this regulation must act upstream of gene expression (reviewed in Ismail *et al*., 2014). When a plant cell is challenged by salinity, it will face a loss of turgor, and, simultaneously, the penetration of sodium through Non-Selective Cation Channels (NSCCs). The turgor loss will activate mechanosensitive calcium channels, culminating in a rapid influx of calcium ions. This will activate the SOS3, a calcineurin-B like sensor and induce its binding to the protein kinase SOS2, resulting in the phosphorylation of SOS1, which will then export the invading sodium ions, driven by a proton antiport from the acidic apoplast to the more neutral cytoplasm (Quintero *et al*., 2002, 2011; Martínez-Atienza *et al*., 2007). The calcium influx will activate NADPH oxidases of the Respiratory burst oxidase Homologue family (Ogasawara *et al*., 2008; Drerup *et al*., 2013), causing an increase of Reactive Oxygen Species (ROS) in the apoplast, which can penetrate the cytoplasm through aquaporins (Bienert *et al*., 2007; Rodrigues *et al*., 2017) and activate a MAP kinase cascade that conveys the signal to the nucleus, turning on transcription factors that deploy a response of gene expression (Xiong and Yang, 2003). The accumulation of calcium and of ROS in the cytoplasm will also activate the biosynthesis of jasmonates (JA), as well as of abscisic acid (ABA) (Hazman *et al*., 2015). The relative timing of these events is decisive for the cellular outcome. A rapid and transient JA signature is linked to cellular adaptation (for instance by boosting the expression of transporters and of enzymatic antioxidants), while a constitutively elevated JA level will culminate in necrosis (Ndecky *et al*., 2023). While necrosis is destructive for the cell itself, it can be an adaptive strategy for the plant as an entity, because older cells act as pools for sodium and divert it from the meristematic cells that can regenerate the plant.

Ionic imbalance will impair stomatal function with negative consequences for photosynthetic efficiency and carbon fixation (for review see Parida and Das, 2005). Stomatal closure can be an indirect effect of turgor loss in the root, releasing signals to the shoot that are composed of physical (such as hydraulic) and chemical (ROS, ABA, possibly peptides) signals (for review see Gorgues *et al*., 2022). However, stomata can also directly respond to the sodium-induced depletion of potassium in the leaf, which will directly impair the ionic fluxes required for opening the guard cells (for review see Shabala and Cuin, 2008). Thus, adaptation to salinity requires cellular as well as systemic events. Therefore, it is evident that it must be orchestrated by phytohormones (for review see Deinlein *et al*., 2014).

In fact, jasmonates (JA) in concert with abscisic acid (ABA) have been identified as crucial determinants of salt susceptibility and tolerance. Rice mutants with a loss of function for the jasmonate biosynthesis gene *Allene Oxide Cyclase* (*AOC*) were found to be more tolerant to salinity (Hazman *et al*., 2015). On the other hand, using a genome-edited mutation of the AOC gene, where the responses of hormones and gene expression were followed over time (Ndecky *et al*., 2023), jasmonates were found to be required for proper water management in the leaf, while being detrimental for sodium retention in the root. This difference was depending on time and organ, indicating that temporal patterns are relevant. A negative feedback loop of the active signal, JA-Ile on the sensitivity to this signal seems to be important, here. This loop derives from the fact that JA-Ile induces the proteolytic decay of JAZ proteins, repressors for jasmonate-inducible genes, whereby the JAZ genes themselves are among the first genes that respond to this de-repression. Thus, jasmonate signalling will rapidly deploy self-inhibition of this signalling. As a result, a transient signature of jasmonate activity emerges that seems central for successful adaptation. In contrast, a constitutive activation of jasmonate signalling is a hallmark for necrosis (for review see Ismail *et al*., 2014). In fact, engineering salt-inducible variants of JAZ proteins that were recalcitrant to proteolysis (evading a constitutive jasmonate signature), was able to cause salt tolerance in rice (Peethambaran *et al*., 2018). Jasmonates activate ABA synthesis and signalling in the leaves. In fact, the leaves of the *aoc* mutant exhibit an impaired salinity response of both ABA-synthesis genes as well as of ABA accumulation (Ndecky *et al*., 2023), contrasting with the roots, where ABA seems to be directly activated by the osmotic component of salinity stress and, therefore, persists in the *aoc* mutant.

Stress-dependent modification of gene expression can either be the consequence of cellular disbalance and, thus, actually a readout of stress damage. They can, however, also reflect events linked to adaptation. To conclude this from the expression pattern *per se* is principally not possible. To discern stress damage from stress adaptation requires additional information, either from a comparison of genotypes contrasting in tolerance, or from dissection of temporal sequences. In order to integrate the regulatory networks inferred from transcriptomics (Ndecky *et al*., 2023), the current study used the same set-up, but focussing on time courses and physiological networks. In particular, ion distribution was followed over time in roots and different parts of the shoot, accompanied by a detailed analysis of biomass allocation between different organs, and a more detailed analysis of photosynthetic parameters beyond stomatal conductance. By supplementing these whole-plant physiological data with targeted RT-qPCR of ion transporters and antioxidant genes over time, we test, which of these responses precede and which follow alterations in physiology. This integrated approach provides the mechanistic resolution necessary to disentangle adaptive from maladaptive contributions of jasmonate signalling to the rice salt stress response and highlights the role of the high-affinity potassium transporter OsHAK16 for successful adaptation, while activation of OsJAZ13 and of the sodium proton antiporter OsNHX1 are identified as hallmarks for salinity-induced damage.

## Materials and methods

### Plant material and growth conditions

The current study compared wild-type (WT) rice (*Oryza sativa* L. *ssp. japonica* cv. Kitaake) and a loss-of-function mutant in *Allene Oxide Cyclase* (*AOC*) generated by CRISPR-Cas9 (Nguyen *et al*., 2020; Ndecky *et al*., 2023). This enzyme is required to generate the jasmonate precursor 12-oxo-phytodienoic acid (OPDA).

Hence, the *aoc* mutant is unable to form jasmonic acid (JA). Since JA depletion leads to male sterility; the homozygous *aoc* mutant seeds had to be isolated from the segregating population of heterozygous *AOC/aoc* plants. The homozygous mutant seedlings can be easily discriminated from heterozygotes or homozygous wildtype by their elongated mesocotyl. To obtain those seedlings, caryopses were surface sterilised by immersion in 70% ethanol (Sigma-Aldrich) for 1 minute, followed by two washes with distilled water. Next, they were soaked in 3% sodium hypochlorite (Sigma-Aldrich) for 20 minutes. Finally, the caryopses were rinsed four times with sterile distilled water to remove residual sterilising agents, and then sown on Murashige and Skoog (MS) medium (8 g.L ¹), solidified with 0.5% phytoagar (Duchefa, Netherlands), in Magenta boxes (Sigma-Aldrich) at 28°C under a 12-h light/12-h dark cycle (Photosynthetically Available Radiation, PAR, ∼475 µmol m ² s ¹) and 60% relative humidity for 6 days. The seedlings were selected for size and homogenous development and then transferred to a hydroponic setup prepared with custom-made, sterilised floating racks (Sigma-Aldrich) placed in glass jars containing Milli-Q water supplemented with 0.358 g.L ¹ of MS basal salts and 1.0 g.L ¹ of MES, adjusted to pH 5.8 (Duchefa, Netherlands). A volume of 500 mL nutrient solution per jar accommodated 20 seedlings, providing 25 mL per individual. Cultivation of the seedlings was continued under the same conditions of light and temperature for additional 11 days. Then, the nutrient solution was replaced with fresh MS medium and either MES as control, or MS medium supplemented with 100 mM NaCl (Sigma-Aldrich) to impose salt stress. To maintain the pH and to prevent growth of cyanobacteria, both solutions were renewed every 3 days. Phenotypic and physiological studies were conducted after 12 days of salt treatment (or transfer to the salt-free medium, respectively) with three biological replicates per condition, each replicate comprising ten seedlings. For molecular studies, the second leaf and roots were collected at four time points: immediately before salt application (0 h) and at 1, 6, and 72 hours after salt application (100 mM NaCl) or transfer to the salt-free medium (control), respectively. Here, each time point and treatment involved three biological replicates consisting of pooled tissue from three seedlings. RT-qPCR was performed using three technical replicates per biological sample. Samples from both control and salt-stressed plants were harvested, immediately frozen in liquid nitrogen, and stored at -80°C until RNA extraction.

### Measurement of growth and fresh weight

Shoot and root lengths were recorded 12 days after salt treatment. Shoot length was defined as the distance from the base of the coleoptile to the tip of the youngest leaf, and root length as the distance from the crown to the tip of the seminal root. For each biological replicate, five individual plants per treatment and genotype were measured. Representative morphological images of WT and *aoc* plants were selected to illustrate phenotypic differences. Growth stimulation by the mutation and growth inhibition by salt were calculated separately for shoots and roots as % change. Growth stimulation by the mutation was calculated within each treatment as (L *aoc* − L*WT*)/L*WT* × 100; growth inhibition by salt was calculated within each genotype as (L *control* − L*salt*)/L *control* × 100, where L is the shoot or root length of the respective group. For the analysis of biomass, the plants were separated into seminal root, the individual leaves 1–5, and the stem including the youngest leaf, which could not be separated from the stem. Fresh weight was measured immediately after harvest and dissection. Data were collected from three biological replicates per genotype and condition, each replicate pooled from ten plants.

### Measurement of photosynthetic parameters

Photosynthetic parameters were measured using a portable photosynthesis system (LI-6800, LI-COR, Lincoln, NE, USA) equipped with a red/blue LED light source under standardised conditions (PAR ∼475 µmol m ² s ¹, relative humidity 60%, temperature 28°C) on the youngest fully expanded leaf (leaf 5) at day 12 after the onset of stress (or transfer to the salt-free medium, respectively). Assimilation rate [µmol.m^-2^s^-1^], transpiration rate [mmol.m^-2^s^-1^], stomatal conductance [mol m ² s ¹], and the efficiency of open photosystem II reaction centres [Fv′/Fm′] were recorded, while PAR was stepwise increased from 0 to 2000 µmol m ² s ¹. Simultaneously, chlorophyll fluorescence was recorded using the LI-6800 fluorometer module, and Fv′/Fm′ was automatically calculated after saturating light pulses. Measurements were collected from both WT and mutant plants for both control and stress conditions, respectively. The measurements were conducted in the morning (in a time window between one and seven hours after dawn). Since all plants needed to be measured sequentially, meaning that the last plant was recorded later than the first plant, the sequence of measurements was rotated to rule out any possible bias from potential diurnal rhythms, such that the values obtained for a given genotype are averaged from different time points during this time window. No systematic differences were detected. Data represent mean and standard error from five weekly replicates.

### Ion content analysis

The individual plant organs were separated as described above, and the harvested tissue dried to complete removal of water at 60°C for 48 h. Then, the dehydrated material was ground into a powder using a pestle and mortar. For the elemental analysis, aliquots of approximately 50 mg of powdered tissue were digested in 2 mL of heated, but not boiling nitric acid (65% v/v), and 0.5 mL of hydrogen peroxide (30% v/v) within closed polypropylene digestion tubes (Gerhardt, UK). The tubes were then heated at 110°C for three hours in a heating block (DigiPrep jr, S-prep). After digestion was completed, the samples were allowed to cool before adjusting the volume to 20 mL with ultra-pure water. The concentrations of sodium and potassium in the samples were determined using the inductively coupled plasma–optical emission spectrometer (ICP–OES, iCAP 7000, Thermo Fisher Scientific). The assay was calibrated using dilutions of standard solutions. Accuracy and precision were checked using certified reference materials digested in parallel with the samples in each batch. For tomato leaves (NIST 1573a), mean recoveries were 100.3% for Na and 91.4% for K, with relative standard deviations of 6.1% and 1.1%, respectively (n = 3 digestion batches). The results are presented as sodium and potassium in mg.g ¹ dry weight, and molar K /Na ratios for each sample. Data represent means and standard errors of three biological replicates.

### Gene expression analysis

The second leaves and roots of the seedlings were excised at specific time intervals (prior to stress, and at 1, 6, and 72 h after the addition of 100 mM NaCl, respectively). At each time point, second leaves from three seedlings were pooled into one sample, and roots from the same seedlings were pooled into a separate sample; three such samples were collected per genotype, treatment and time point. The harvested tissues were immediately frozen in liquid nitrogen and stored at -80°C until use. Total RNA was extracted using the innuPrep plant RNA kit (Analytik Jena) according to the protocol of the manufacturer. Quantity, quality, and integrity of the RNA were verified spectrophotometrically (NanoDrop) as well as by agarose gel electrophoresis. Reverse transcription and quantitative real-time PCR were conducted as described by Hazman *et al*. (2015). The amplification specificity was confirmed by melt-curve analysis. Gene expression was normalised to UBQ10 as reference gene, and relative transcript levels calculated using the 2 ^ΔCt^ method (Livak and Schmittgen, 2001). Raw Ct values of UBQ10 were stable across treatments and time points. Data represent mean and standard error from three biological replicates, each in technical triplicates. The details of the oligonucleotide primers to amplify the genes of interest are provided in **Supplementary Table S1**.

### Statistical analysis

Shoot and root length were determined on three biological replicates per genotype and condition, each based on five individual plants. Fresh weight and ion content were determined on three biological replicates, each pooled from ten plants. Photosynthetic parameters were determined in four independent experiments performed in consecutive weeks. Gene expression was determined on three biological replicates, each pooled from three seedlings, with three technical replicates per biological sample. Data were expressed as mean ± standard error (SE). Statistical significance between wild type and *aoc* was determined separately for each organ, treatment and time point using two-sided unpaired Student’s t-test. Differences were considered significant at \**P* < 0.05, \*\**P* < 0.01 and \*\*\**P* < 0.001.

## Results

### The *aoc* mutant performs better under salinity stress

To understand the role of jasmonates on growth responses to salinity, we compared the *japonica* cultivar Kitaake (wild type, WT) and a mutant, where *Allene Oxide Cyclase* (*AOC*), a key enzyme for the biosynthesis of jasmonic acid, had been inactivated by CRISPR-Cas9 (Nguyen *et al*., 2020). Since AOC is a single-copy gene in rice, this mutant is not able to accumulate jasmonates. The phenotypic difference between the two lines that differ exclusively with respect to the activity of the *AOC* gene was assessed at day 12 after transfer to the hydroponic set-up, either in the absence (control) or presence of 100 mM NaCl. The salt effect on the morphology was distinct in both genotypes. Plantlets of the WT were shorter, more chlorotic and necrotic than those of the *aoc* mutant, which displayed a greener foliage (**Fig. 1A, B**). However, there was also a clear effect of genotype, since the mutant was bigger even under control conditions. When we quantified the growth of shoots and roots, we found that the shoot was significantly longer than WT under both, non-saline and saline conditions (**Fig. 1C**). Also for the root, length was increased in the mutant, for both conditions (**Fig. 1D**). To separate the effects of the genotype from those of salinity, we also plotted the relative changes, once for the genotype (**Fig. 1E**), once for the effect of salt stress (**Fig. 1F**). This approach revealed that the shoot in the mutant is about 20% longer, irrespective of the stress condition, while for the root, the enhanced growth drops from 55% under control conditions to 25% under salinity (**Fig. 1E**). The relative inhibition of growth by salinity was comparable between WT and mutant for the shoot (around 25%), while for the root it was less in the WT, with around 10% as compared to around 25% in the mutant (**Fig. 1F**). In summary, the *aoc* mutant can sustain a greener appearance under salinity, and the same trend was reflected in the biomass data (**Fig. 2**). In addition, the growth benefit from the mutation is seen also under salinity, mainly for the shoot, less pronounced for the root. It should be noted that the superior performance of the *aoc* mutant is seen already in the absence of salinity.

**Fig. 1.**
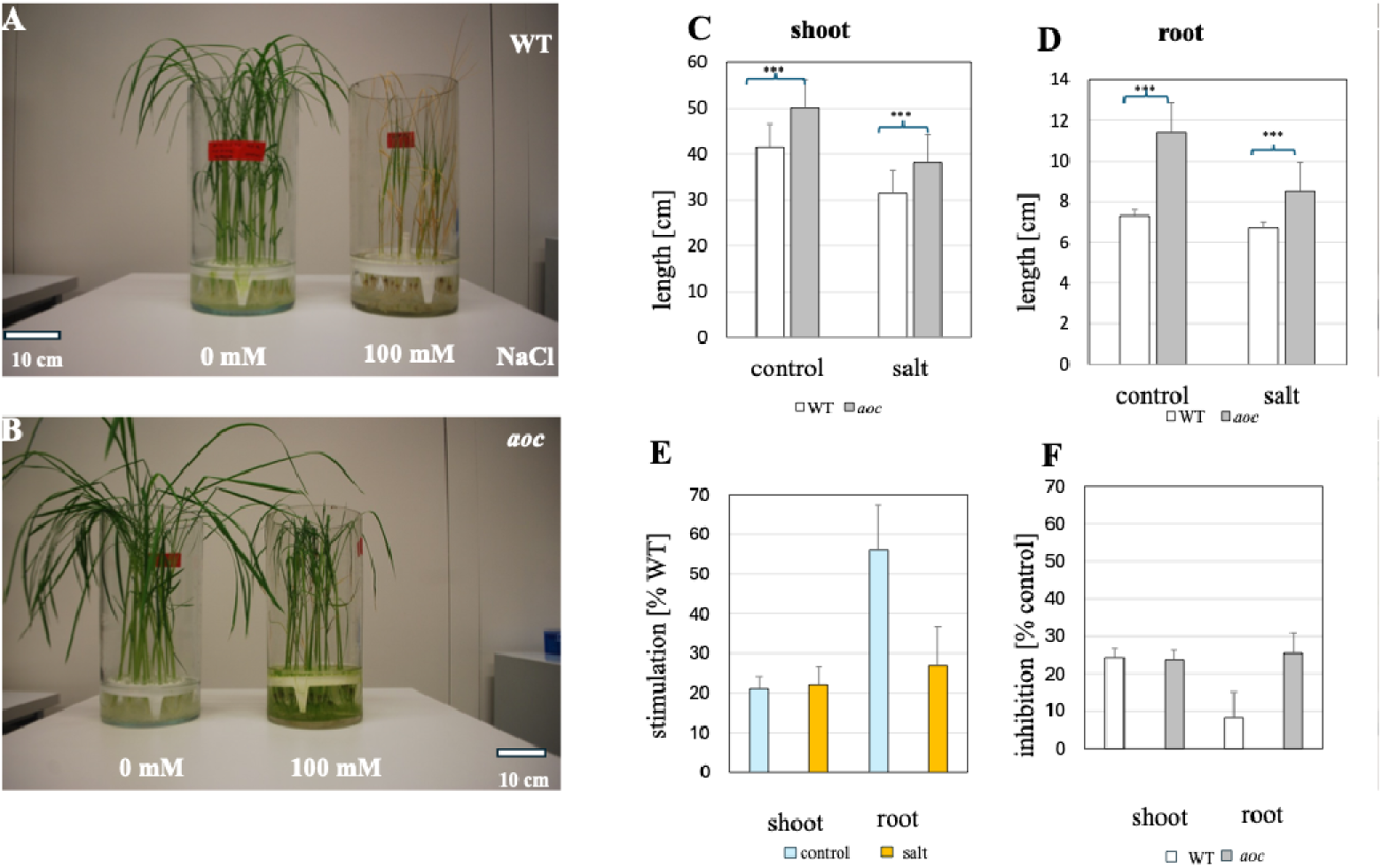
Phenotypical effect of the CRISPR-Cas9 generated *aoc* null mutation in the background of the *ssp. japonica* cultivar Kitaake after 12 d of salinity stress. Representative individuals are shown for the WT (**A**) and the *aoc* mutant (**B**). Length of shoot (**C**) and root (**D**) at the end of the stress treatment. Data represent mean and standard error of three biological replicates, each based on five individual plants. (**E)** Dependence of growth stimulation in the *aoc* mutant on type of organ and stress condition. Note that the stimulation in the shoot is independent of the stress condition, while in the root, it is strongly dependent. (**F)** Dependence of salt induced growth inhibition on type of organ and genotype. Note that the inhibition by salt is the same with exception of roots of the WT, that are less inhibited by salt as compared to the roots of the *aoc* mutant.

**Fig. 2.**
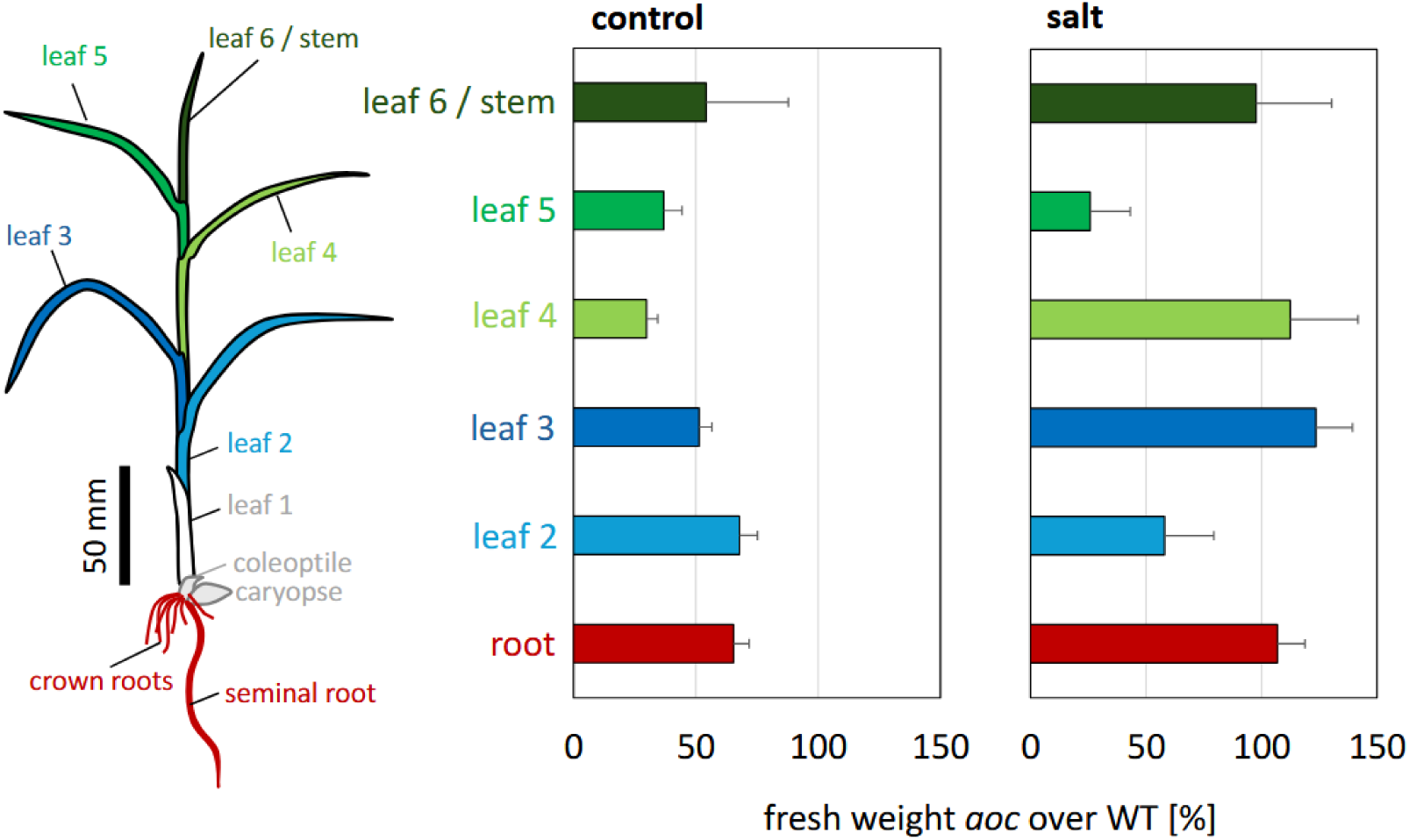
Developmental dependence of the mutant effect based on fresh weight. The growth stimulation in the *aoc* mutant over the WT, calculated as (FW*aoc* − FWWT)/FWWT × 100, is plotted for the different organs of the plant either in control conditions or after 12 d of salinity stress (100 mM NaCl). Data represent mean and standard error of three biological replicates, each comprising ten plants. For each leaf, blade and sheath were collected together. The youngest leaf was not expanded yet and, therefore, could not be separated from the subsequent leaves and the meristem.

### Salinity increases fresh weight in *aoc* leaves through cell expansion

To get insight into the cellular mechanisms behind the enhanced growth of the mutant, we measured the fresh weight of individual organs (**Fig. 2**). Here, the jasmonic acid deficient *aoc* mutant accumulated more fresh weight than the wild type across all organs. This difference was already seen under control conditions, but became accentuated under salinity, meaning that the mutant organs retained their growth more efficiently than the wild type. This effect was particularly evident in the younger leaves 3 to 4 (also in the part comprising leaf 6 and stem), whereas the more mature leaf 2 displayed only minor differences between genotypes. Likewise, the root of the mutant preserved its fresh weight more efficiently than the wild type. A difference in fresh weight could be either due to an increase in cell number, or to an increase in cell volume. In the second case, dry weight should behave in a manner parallel to fresh weight, in the first case, they should diverge. When we compared the patterns for fresh weight and dry weight (**Suppl. Fig. S1**), we observed, for the root, that the salinity response of the dry-weight surplus of the mutant increased with a similar factor as seen for the fresh-weight surplus, suggesting that this phenomenon was based on cell number, i.e. cell proliferation. This proportional scaling was very similar for control and saline conditions. The situation was qualitatively different for the response of leaves 3 and 4 (**Suppl. Fig. S1**, red arrows). Here, the salient fresh-weight surplus of the mutant seen under salinity was not accompanied by a corresponding increase of dry weight. This means that this increase of fresh weight is not due to increased cell proliferation, but of cell expansion. In summary, these data reveal that the *aoc* mutant sustains growth under salinity through organ-specific mechanisms promoting cellular expansion in the shoot while maintaining proliferative capacity in the root.

### The *aoc* mutant sustains photosynthetic performance more efficiently under salinity

To get insight into the role of jasmonates for the photosynthetic response to salinity, we followed several photosynthetic parameters in wild type and *aoc* mutant (**Fig. 3**). Salinity negatively affected carbon assimilation in both lines, however, to a different extent (**Fig. 3A**). When the fluence dependency of assimilation rate was recorded, the mutant showed saturation at around 30% lower rates than the WT. Under salinity, the relationship was inverted. Here, the WT dropped to almost zero, while the *aoc* mutant still sustained around 30% of the value seen under control conditions. For both transpiration rate (**Fig. 3B**) and stomatal conductance (**Fig. 3C**), the mutant displayed, under control conditions, higher values (around 50%) as compared to the WT. However, while both parameters dropped to zero in the wild type in response to salinity, the mutant was still able to produce around 25% of transpiration and stomatal conductance compared to the values seen in the absence of salinity. Interestingly, the efficiency of open photosystem II reaction centres (Fv′/Fm′) was almost the same for WT and *aoc* mutant, no matter, whether assessed under control conditions, or under salinity (**Fig. 3D**). Here, the curves also converged at similar values, when control and stress conditions were compared.

**Figure 3.**
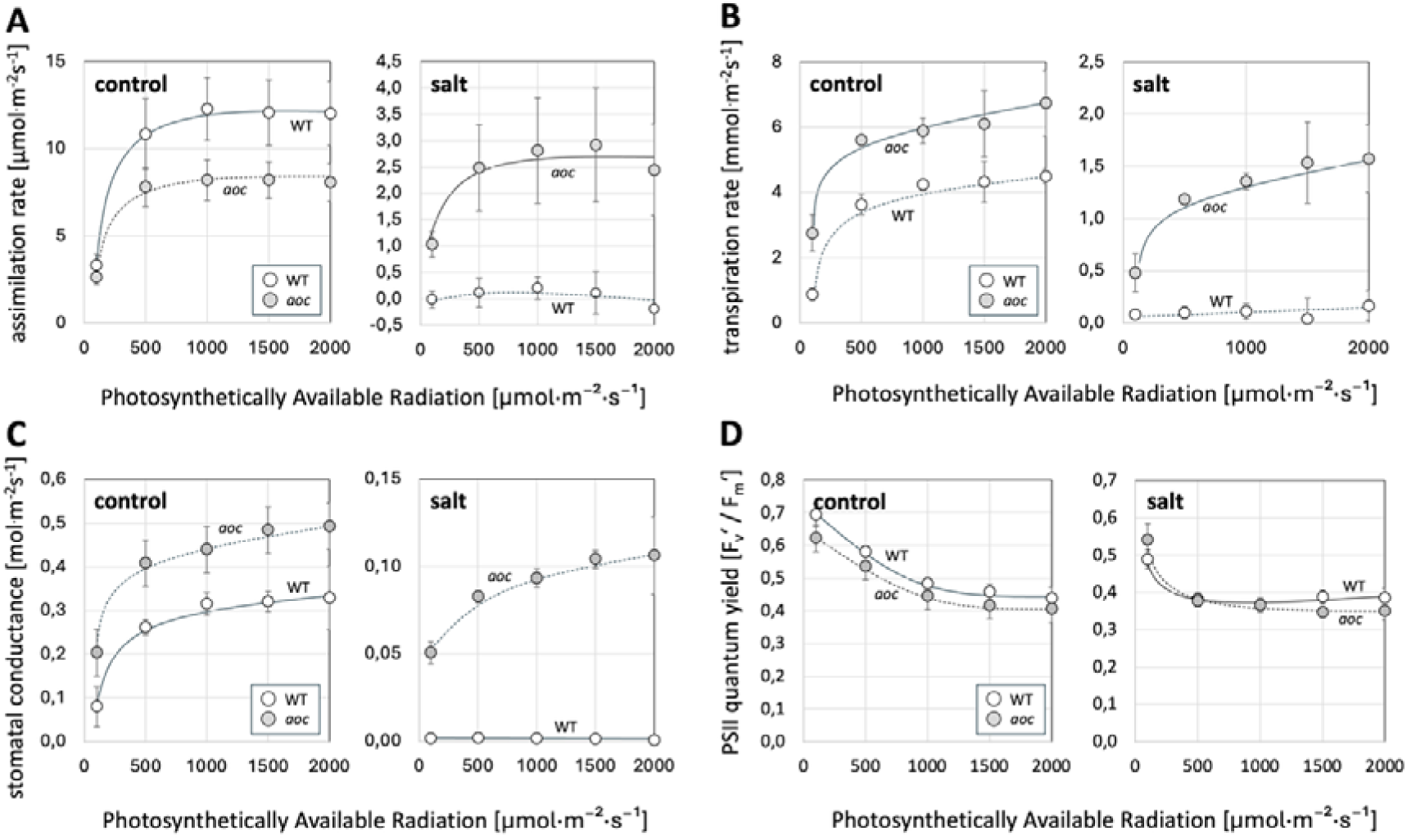
Dependence of photosynthetic parameters on light intensity in light-adapted plants of the WT versus the *aoc* mutant. (**A)** assimilation rate (**B)** transpiration rate (**C)** stomatal conductance (**D)** efficiency of open photosystem II reaction centres (Fv′/Fm′). Measurements were performed on the fifth leaf after 12 days of salt treatment (100 mM NaCl) or transfer to salt-free medium (control), across a range of photosynthetically available radiation (PAR, 0– 2000 µmol m ² s ¹). Data represent means and standard errors of four independent experiments.

In summary, the jasmonate-free *aoc* mutant can cope more efficiently with the challenges of salinity as compared to the WT, consistent with the enhanced growth and biomass accumulation observed under salinity. The superior performance of assimilation rate is correlated with the retention of stomatal conductance and transpiration in the mutant, whereas both parameters declined to near-zero values in the WT . In contrast, Fv′/Fm′ is not revealing any significant differences between WT and mutant.

### The *aoc* mutant partitions sodium and potassium differently

To link the physiological differences between WT and *aoc* mutant with differences in ion homeostasis, we monitored sodium content across the different organs. Under control conditions, the sodium levels in the mutant were conspicuously lower than in the WT across all organs, with the exception of leaf 5 (Fig. 4). However, in response to salt stress, both genotypes showed a high accumulation (in the range of two orders of magnitude across the control levels) of sodium in both, the shoot and the root. For the WT, this was especially conspicuous in leaves 2, 4, and 6 (**Fig. 4**). For the mutant, uptake into the aerial tissues was strongly reduced, prominently in leaves 2, 4, and 6 (around 25-30% lower than in the WT), confirming the reduced sodium accumulation reported for the second leaf by Ndecky et al. (2023). Interestingly, this reduced uptake was not seen in the root, where sodium levels under salinity did not differ between WT and mutant. The increase in sodium was accompanied by a sharp drop in potassium levels (**Suppl. Fig. S2**). This drop was especially pronounced for the youngest parts of the shoot (leaf 6 / stem), where the values dropped 4 to 5-fold as compared to the control. Interestingly, the resting levels of potassium were consistently higher in the mutant throughout all organs. Under salinity, this surplus disappeared completely. For leaves 3 and 4, the potassium levels under salinity were even lower than in the WT. When the molar K /Na ratio was determined (**Fig. 5**), the mutant showed clearly elevated values over all organs, which is due to the lower resting levels of sodium (**Fig. 4**) and the higher resting levels of potassium (**Suppl. Fig. S2**). Under salinity, this ratio plummeted by almost three orders of magnitude, albeit with organ dependent differences. For leaves 5 and 2, the ratios tended to be higher in the mutant, for leaves 3 and 4, the ratios were higher in the WT. Overall, the *aoc* mutant can exclude sodium more efficiently, especially from young leaves.

**Figure 4.**
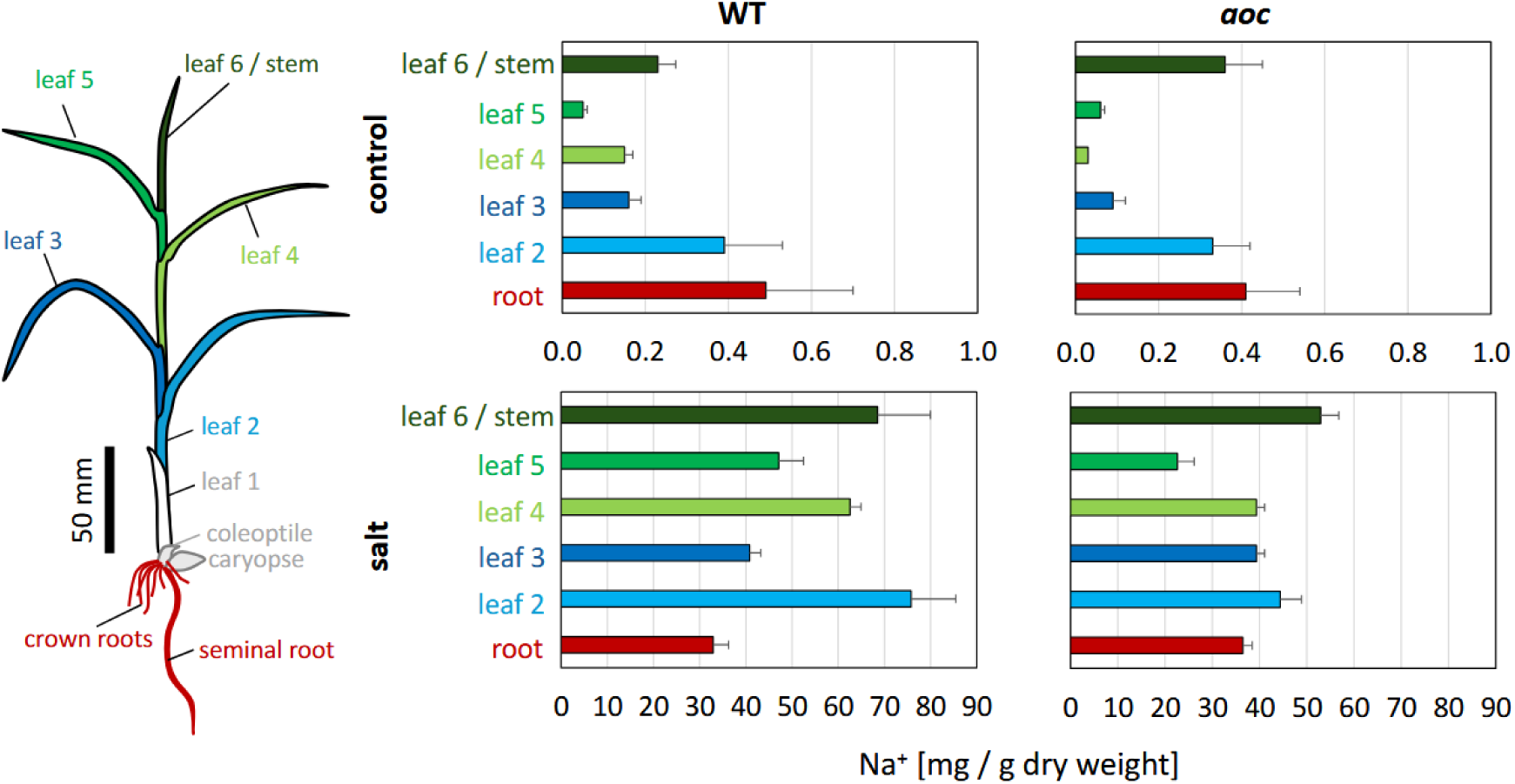
Organ-specific differences in sodium accumulation between the WT and the *aoc* mutant. Sodium contents in WT and *aoc* mutant are plotted for the different organs of the plant either in control conditions or after 12 d of salinity stress (100 mM NaCl). Data represent mean and standard error of three biological replicates, each comprising ten plants. For each leaf, blade, collar, and leaf sheath were collected. The youngest leaf was not expanded yet and, therefore, could not be separated from the subsequent leaves and the meristem.

**Figure 5.**
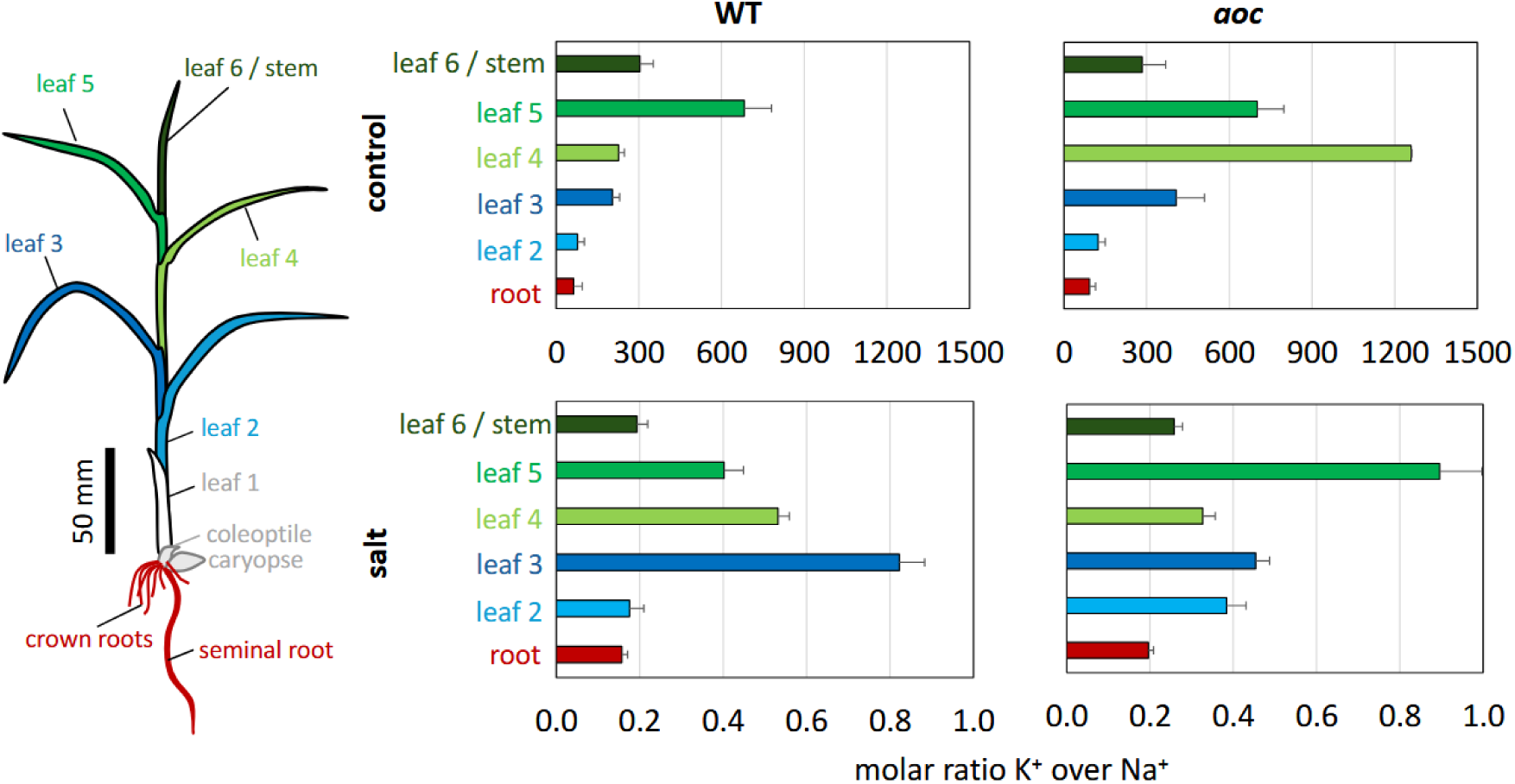
Organ-specific differences in the potassium/sodium ratio between the WT and the *aoc* mutant. Potassium/sodium ratios are plotted for the different organs of the plant either in control conditions or after 12 d of salinity stress (100 mM NaCl). Data represent mean and standard error of three biological replicates, each comprising ten plants. For each leaf, blade, collar, and leaf sheath were collected. The youngest leaf was not expanded yet and, therefore, could not be separated from the subsequent leaves and the meristem.

### Salinity induces *JAZ* genes depending on jasmonates

To examine how salinity modulates jasmonate biosynthesis and signalling, steady-state transcript levels of *JAR1* (conjugating the inactive precursor jasmonic acid to isoleucine, leading to the bioactive JA-Ile) and a set of *JAZ* genes were followed over time in leaf tissue of WT and *aoc* mutant plants kept either under control conditions or salinity treatment (**Fig. 6**). Transcripts for JAR1 showed a slight transient decrease for the WT under salinity (Fig. 6A). For the mutant, there was a slight transient increase under the same conditions. Under control conditions, there was a pronounced increase by a factor of four in the mutant, which was not seen in the WT, but already at 6 h, transcript levels had returned to the resting levels. All four tested *JAZ* transcripts *JAZ8*, *JAZ9*, *JAZ11*, and *JAZ13* were up-regulated by salinity in the WT, while expression in the *aoc* mutant was almost non-detectable (**Figs. 6B–E**). The amplitude and the timing of induction varied: *JAZ9* and *JAZ11* increased steadily from the first time point (1 h), but at a moderate slope. In contrast, *JAZ13* accumulated with about three times the rate, also detectable from 1 h. The behaviour of JAZ8 was more complex, a small early and transient increase was followed by a later and more pronounced accumulation detected at 72 h. Thus, *JAZ13* was found to be the most salt-responsive of the tested *JAZ* transcripts. The complete absence of any induction in case of the mutant indicates that the salinity-induced expression of all tested JAZ genes strictly requires jasmonates.

**Figure 6.**
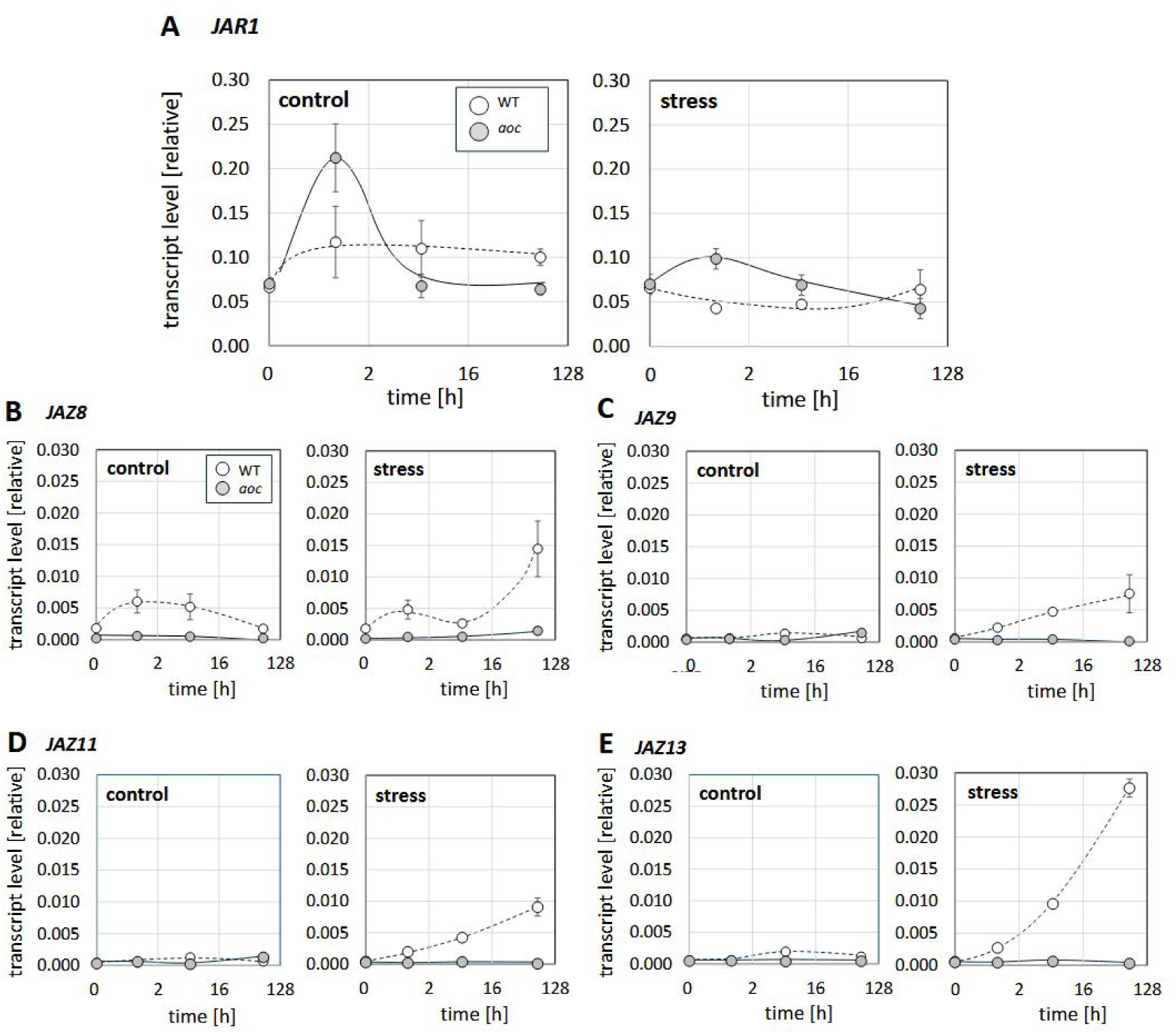
Steady-state transcript levels in leaf 2 for a panel of jasmonate-related genes in WT and *aoc* mutant either in control conditions or under salinity stress (100 mM NaCl). The probed genes were the jasmonic acid isoleucine ligase (*JAR1,* **A**), and the JAZ repressors *JAZ8* (**B**), *JAZ9* (**C**)*, JAZ11* (**D**), and *JAZ13* (**E**) known as salt responsive. Data represent mean and standard errors from three biological replicates, each in technical triplicates.

### The salinity response of *HAK16* is specifically enhanced in the *aoc* mutant

The reduced accumulation of sodium into leaf 2 was among the most salient differences in ion distribution between WT and mutant (**Fig. 4**). To get the context of this phenomenon, time courses for the expression of the vacuolar sodium transporter *NHX1* (**Fig. 7A**), and three high-affinity potassium transporters (**Fig. 7B-D**) were followed in leaf 2, either under control conditions, or under salinity. These transporters were selected because they act at two decisive steps determining leaf sodium levels: vacuolar sequestration of sodium (*NHX1*) and long-distance potassium transport (*HAK* family), which shapes the potassium/sodium balance of the leaf.

**Figure 7.**
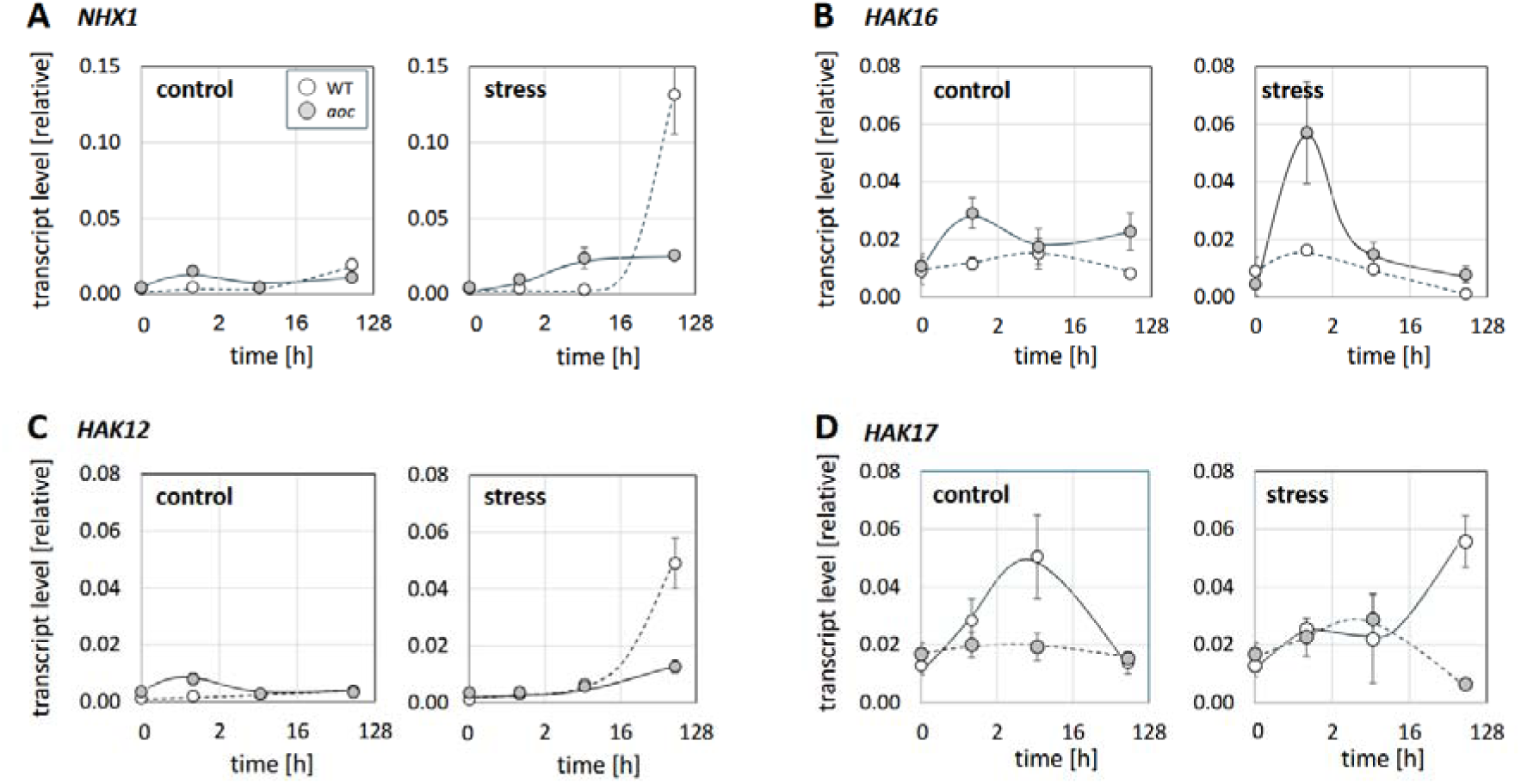
Steady-state transcript levels in leaf 2 for a panel of ion transporters in WT and *aoc* mutant either in control conditions or under salinity stress (100 mM NaCl). Data represent mean and standard error of three biological replicates, each in technical triplicates. The probed genes were the vacuolar sodium-proton antiporter (*NHX1,* **A**), the plasma membrane located high-affinity potassium (*HAK*) transporters *HAK12* (proposed to exclude sodium from the shoot, **C**), *HAK16* (proposed to drive potassium uptake in the root and translocation to the shoot, **B**), and *HAK17* (proposed to mediate potassium uptake under high sodium concentrations, **D**).

The steady-state levels of *NHX1* transcripts remained stable and comparable between WT and *aoc* in the absence of salinity. Upon salt stress, this vacuolar sodium antiporter was induced in the mutant already at 1 h, while the WT did not respond at all. Only for the last time point, 72 h, the pattern was reversed. Here, the WT displayed very high transcript levels exceeding those in the mutant by around 5-fold. For the high-affinity potassium transporters *HAK12*, discussed to extrude sodium from the shoot (**Fig. 7C**), transcript levels remained low and constant in both genotypes under control conditions, while under salinity the WT showed a pronounced induction at 72 h that was absent in the mutant. In the mutant, only a small transient rise was observed at 1 h under control conditions. For *HAK16*, thought to mediate potassium uptake in the root and translocation to the shoot (**Fig. 7B**), the pattern was opposite to that seen for *NHX1* (**Fig. 7A**), i.e., a strong and **early** induction under salinity **in the mutant**, peaking around 1–2 h, absent from the WT. For *HAK17*, mediating potassium uptake under the pressure of high sodium (**Fig. 7D**), there were no salient responses for the mutant, while the WT showed a transient (control) or a late (salinity) induction. Thus, the elevated accumulation of sodium in leaf 2 of the WT correlates with the strong expression of NHX1 at 72 h (Fig. 7A), which is accompanied by elevated expression of *HAK12* and *HAK17* under the same conditions. By their temporal signatures, these responses need to be seen as damage responses to the elevated accumulation of sodium. Consistently, these inductions are not observed in the *aoc* mutant, where sodium accumulation into leaf 2 is substantially weaker as in the WT.

The reduced sodium accumulation into leaf 2 of the mutant might be linked with the more pronounced sequestration of sodium (**Fig. 4**) and potassium (**Suppl. Fig. S2**) in the root. Therefore, we analysed transcripts for the same panel of ion transporters (**Suppl. Fig. S3**). However, the vacuolar sodium–proton antiporter NHX1 (**Suppl. Fig. S3A**) remained uninduced and comparable between WT and *aoc* mutant plants under both control and salinity conditions.

In contrast, the pattern for *HAK16*, the transporter responsible for the uptake of potassium into the root and its translocation into the shoot, contrasted clearly between the two genotypes (**Suppl. Fig. S3B**). While there was no significant difference under control conditions, these transcripts were steadily increasing in the mutant, while they did not respond in the WT. However, this induction in the mutant was definitely lagging behind the rapid and fast induction seen in the leaves (**Fig. 7B**). In contrast, *HAK12* (**Suppl. Fig. 3C**), while induced by salinity, did not exhibit a difference between mutant and WT; and for *HAK17* there was no significant difference, neither between WT and mutant, nor between control and salinity (**Suppl. Fig. 3D**). Overall, transcriptional responses of ion transporters in the root were limited and delayed compared to those observed in the shoot. This indicates that the major differences in sodium exclusion and potassium partitioning between WT and *aoc* mutant plants are predominantly established in the shoot rather than in the root, consistent with the transcriptome analysis of Ndecky et al. (2023), where more transporter genes were found to be modulated in a jasmonate-dependent manner in the shoot than in the root.

### The *aoc* mutant shows elevated expression of plastidic Superoxide Dismutase

Sodium ions interfere with electron transport at the thylakoid resulting in elevated production of superoxide. Superoxide needs to be detoxified by Superoxide Dismutase (SOD) into hydrogen peroxide, which is then further dissipated by catalases and peroxidases. We assessed, therefore, the transcrip Ds (**Fig. 8**). The expression of both genes, *ptSOD1* and *ptSOD2* increased around fourfold in the *aoc* mutant, peaking at 6 h after the onset of salt stress. This induction was not seen in the WT. It should be noted, however, that the induction of the two *SOD* genes was also observed under control conditions, to a fairly similar extent, indicating that these responses were caused by the transfer to the hydroponic system, rather than by salt stress. Thus, plastidic *SODs* seem to be very sensitive markers to any kind of perturbation, not necessarily specific for salinity. Irrespective of this, these genes seem to be more readily induced in the *aoc* mutant as compared to the WT.

**Figure 8.**
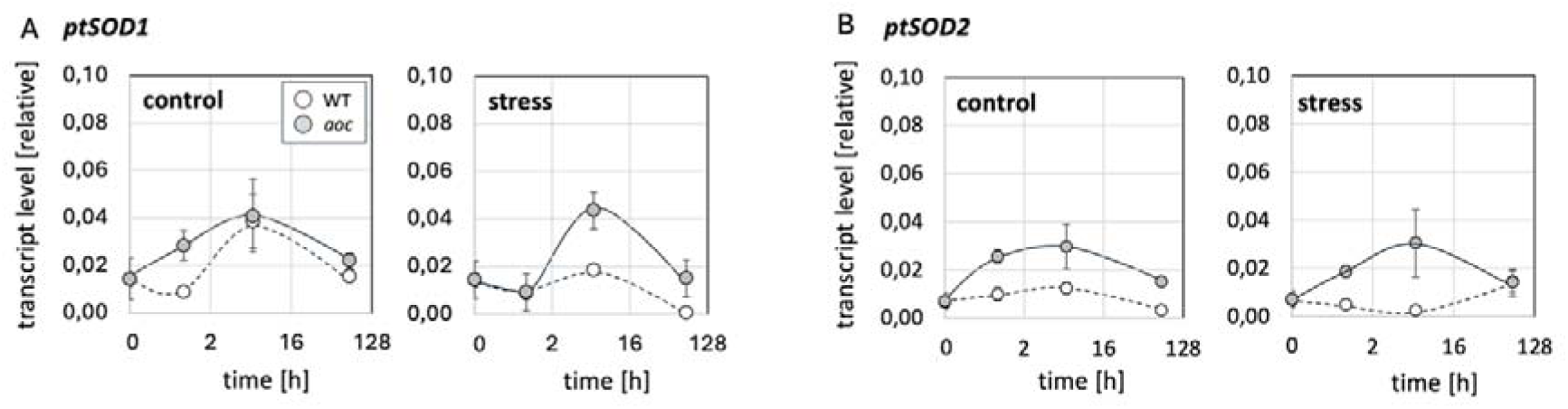
Steady-state transcript levels in leaf 2 for the plastidic superoxide dismutases *ptSOD1* (**A**) and *ptSOD2* (**B**) in WT and *aoc* mutant either in control conditions or under salinity stress (100 mM NaCl). Data represent mean and standard error of three biological replicates, each in technical triplicates.

## Discussion

Jasmonates are central signals coordinating cellular and systemic responses to abiotic stress factors, including osmotic and salinity stress (for review see Riemann *et al*., 2015). For rice, mutants impaired in the synthesis of jasmonic acid, have been found to cope with salinity more efficiently indicating that jasmonates act as negative regulators in this context (Hazman *et al*., 2015). To understand the molecular base of this improved salinity tolerance, we had previously monitored, by RNAseq, differentially expressed transcripts in a CRISPR-Cas9 generated mutant of *Allene Oxide Cyclase* (*AOC*) compared to the wild-type background, the *japonica* cultivar Kitaake (Ndecky *et al*., 2023). The current study extends these transcriptomics and hormone analysis data by the physiological context. We followed different physiological traits, including a non-invasive monitoring of photosynthetic parameters, the distribution of sodium and potassium ions as well as of biomass between different organs of the plant over time and monitored the time courses of transcripts for ion transporters, jasmonate response, and superoxide scavenging, to understand the temporal sequence of these responses to salinity. We find a correlation between a late deregulated activation of the jasmonate pathway, a rapid stomatal closure, and a late activation of the vacuolar antiporter *OsNHX1* with salinity induced damage. In contrast, the early activation of the high-affinity potassium transporter *OsHAK16* correlates with a reduced allocation of sodium to young leaves. These key findings lead to the following questions that are addressed in the following discussion: What is the role of jasmonates for salinity-induced stomatal closure? What can be inferred about the role of *OsHAK16* for adaptation? What is the functional relevance of the late increase in jasmonate activity?

### Does the lack of OPDA help the *aoc* mutant to keep stomata open under salinity?

The inhibited growth in the WT might be caused by salt-induced damage to chloroplasts. However, Fv′/Fm′ remained essentially unchanged and did not differ between the two genotypes (**Fig. 3**). In contrast, stomatal conductance and transpiration were almost completely shut down in the WT, while the mutant retained still 25-30% of conductance. This shifts stomata into the focus. Jasmonates might either act operationally, by supporting a full closure response to salinity. They might, however, also act structurally, by altering stomatal patterning during leaf development.

For the operational hypothesis, the interaction between jasmonates and ABA has to be considered: The transient induction of ABA in the leaves is mildly delayed in the *aoc* mutant, but still comparable, if the integral over time is considered (Fig. 6B in Ndecky *et al*., 2023), indicating that the sustained transpiration cannot be explained in terms of reduced ABA biosynthesis alone. In fact, the induction of ABA-responsive genes such as RAB and dehydrins is strongly impaired in the aoc leaf (Fig. 6C in Ndecky et al., 2023), so that the defect appears to reside downstream of the hormone, in a reduced responsiveness to ABA. In fact, in addition to the canonical drought response mediated by ABA, stomata can also close in response to the jasmonate precursor OPDA (Savchenko *et al*., 2014). Several signalling components seem to be shared by both pathways. For instance, the *abi2-1* mutant, impaired in ABA signalling, cannot close stomata in response to MeJA (Munemasa *et al*., 2007), and a calcium-triggered activation of the CBL1-CIPK5 phosphorylation pathway leading to activation of the potassium efflux channel GORK is not only induced by ABA, but also through jasmonate signalling evoked by localised wounding (Förster *et al*., 2019). Allene Oxide Cyclase is the enzyme generating OPDA. Thus, the *aoc* mutant not only fails to accumulate jasmonate isoleucine, but also its precursor OPDA, which seems to be the actual player in oxylipin dependent stomatal closure (Savchenko *et al*., 2014). Whether the sustained transpiration in the *aoc* mutant is caused on the operational level, remains doubtful, however. At least in Arabidopsis, the stomatal closure induced by ABA in the *coi1* mutant (not able to process the JA-Ile signal) remains comparable to that of the WT (Fig. 1 in Munemasa *et al*., 2007)

This brings the structural scenario into the discussion. In fact, overexpression of *JAZ9* in rice produced to leaves with reduced stomatal density, and, hence, lower transpiration rates (Singh *et al*., 2021). The fact that both, transpiration rates and stomatal conductance are substantially elevated in the *aoc* mutant over the WT already under control conditions (**Fig. 3B, C**) further supports a scenario, where jasmonate signalling acts as negative regulator for either inception or differentiation of guard-cell forming meristemoids. This is supported by observations in Arabidopsis, where methyl jasmonate reduced stomata number, while impaired jasmonate biosynthesis or perception increased stomatal development, acting through MYC2, MYC3 and MYC4 upstream of SPEECHLESS and FAMA (Han *et al*., 2018).

Two implications of the two concurrent models would be directly testable experimentally. Exogenous application of JA-Ile or OPDA should rescue the operational, but not the structural stomata phenotype of the mutant. On the basis of the operational model, the transpiration rate under salinity should be eliminated mimicking the situation in the WT. Genetic interference with a step downstream of OPDA, but upstream of JA (for instance, loss of function of OPR activity, or OPDA transport from plastid to peroxisomes) should bring transpiration under salinity down to the level seen in the WT, while the elevated transpiration rate under control conditions should resemble that of the *aoc* mutant.

### The early induction of *OsHAK16* in roots and leaves is a hallmark of salt adaptation

The capacity of the *aoc* mutant to sustain growth under salinity is also reflected at the level of biomass allocation. The mutant retained fresh weight more efficiently than the wild type across all organs, most notably in the younger leaves 3–4 and in the root (**Fig. 2**). Comparison of fresh and dry weight (**Suppl. Fig. S1**) indicated that this advantage rests on organ-specific mechanisms: in the shoot it reflects cell expansion, whereas in the root the parallel rise in dry weight points to sustained cell proliferation. Maintaining turgor-driven expansion under salinity requires preservation of the K /Na balance (for review see Shabala and Cuin, 2008), meaning that the sustained growth in the mutant must be linked with altered activity patterns of ion transporters.

Among the tested transporters, two exhibited a salient difference in their response. The elevated accumulation of sodium in leaf 2 of the WT correlated with a strong expression of *NHX1*, an induction that was not seen in the *aoc* mutant (**Fig. 7A**). In contrast, for the root, this transporter did not show any response at all, neither in the WT, nor in the *aoc* mutant. This transporter is located in the tonoplast and removes cytosolic sodium ions by sequestration into the vacuole. Since the overexpression of *NHX1* can improve salt tolerance in rice (Fukuda *et al*., 2004), it might be considered as adaptive. However, we did not detect any induction of this transporter in the root, neither in the WT, nor in the mutant (**Suppl. Fig. S3A**), and the induction in the leaf was late, correlating with the local sodium concentration. Thus, in rice, the induction of *NHX1* seems to be a damage response to elevated sodium. This contrasts sharply with the situation in the salt-tolerant crop sorghum, where efficient sequestration of sodium into the distal elongation zone of the root was found to be a hallmark for adaptation (Abuslima *et al*., 2022).

The second transporter candidate was the high-affinity potassium transporter *OsHAK16*, a member of the HAK/KUP/KT family. These transporters are crucial for K /Na homeostasis, preserving membrane potential, potassium-dependent metabolic activity, and stabilising cell expansion, while simultaneously circumventing K -efflux-induced programmed cell death (Shabala and Cuin, 2008). Basically, they convey their function either by retrieval of sodium from the transpiration stream, or by acquisition of potassium from the rhizosphere (reviewed in Horie *et al*., 2009 and Dave *et al*., 2022). For instance, OsHKT1 was identified from a QTL for salinity tolerance and then shown to extract sodium from root stele and basal nodes to protect the youngest photosynthetically active leaves in rice (Ren *et al*., 2005). In contrast, other transporters of this family, such as OsHAK1, OsHAK5, and OsHAK16 exert their effect by promoting K^+^ transfer from the soil into root epidermis and stele (Yang *et al*., 2014; Chen *et al*., 2015; Feng *et al*., 2019). In our study, the *OsHAK16* transcript was found to be rapidly and steadily salt-induced in the root of the *aoc* mutant, but not in the WT (**Suppl. Fig. S3B**). In the leaves, a rapid and transient induction was seen for both genotypes, but at a doubled amplitude in the *aoc* mutant (**Fig. 7B**). This transporter was already identified as salt inducible during previous expression profiling (Okada *et al*., 2008). Also, in our earlier transcriptome analysis (Ndecky *et al*., 2023), *OsHAK16* was among the transporters that, although not being the most abundant, showed the strongest salt induction in the roots of the *aoc* mutant. The functional impact of this transporter is also demonstrated by a study with rice lines where *OsHAK16* was either overexpressed or functionally null by a T-DNA insertion (Feng *et al*., 2019) demonstrated an adaptive role for this member of the HAK family. In contrast to NHX1, this transcript is induced rapidly, qualifying it as a candidate for salinity adaptation, rather than as marker for salinity-inflicted damage. The link of this transporter to jasmonates might be mediated by OsJAZ9, a transcriptional regulator that can repress OsbHLH062, a transcriptional activator that can bind on a motif, termed E-box, in the promoters of HAK transporters (Wu *et al*., 2015). In fact, inactivation of *OsJAZ9* deployed a salinity tolerance that was linked with improved potassium homeostasis. An implication of a scenario, where jasmonate, acts as negative regulator of HAK expression would be that salinity-induced damage in the WT should go along with an induction of *OsJAZ9*, which is, what we could observe (**Fig. 6C**).

Three further predictions of this model can be tested in future experiments. First, introducing a *hak16* knockout in the jasmonate-deficient background should at least partially restore the salt-susceptibility seen in the WT. Second, pre-treatment with MeJA prior to salt stress, should suppress the induction of *HAK16* and eliminate the salt tolerance of the aoc mutant. Third, the higher residual potassium levels found in the *aoc* mutant under salinity (**Suppl. Fig. S2**) should come with a greater net K retention and reduced GORK-type K efflux in direct flux measurements. If so, this would provide a common mechanism for both, the sustained potassium pool in the root and the sustained stomatal aperture discussed above.

### Adaptation versus necrosis - a matter of jasmonate timing?

Jasmonate signalling is endowed with a negative feedback loop leading to a specific temporal signature, pivotal for adaptation to the triggering stress. The active form of jasmonic acid, JA-Ile, promotes COI1- mediated degradation of JAZ repressors via the SCF/COI1 ubiquitin–proteasome pathway. As a result, repression of jasmonate-regulated genes is released. Among the earliest transcripts seen in response to JA-Ile, are the JAZ genes themselves (for review see Wasternack and Song, 2017). In our experiments, *OsJAZ8* (**Fig. 6B**) and, with some delay, but more vigorously, *OsJAZ13* (**Fig. 6E**) responded most rapidly to salinity in the WT, marking a vigorous attempt to close the feedback loop, which acts like a “brake” restraining stress signalling to a temporally defined signature avoiding a constitutive and de-regulated stress status. This biphasic stress response recalls Hans Selye’s (1974) original concept of adaptive “eustress” and destructive “distress”.

However, this early response was followed by a later and massive accumulation at the latest time point (3 d). In case of *OsJAZ8*, this increase was preceded by a transient decrease. Neither the early induction, nor the late accumulation was seen in the *aoc* mutant, which is to be expected from a model, where activation of *JAZ* genes is dependent on JA-Ile. To interpret the biphasic accumulation of *OsJAZ8* one needs to consider the time course of jasmonate abundance, which was investigated during our preceding study (Ndecky *et al*., 2023). Here, for the WT (the *aoc* mutant was not responsive) the precursors OPDA and JA, but also the bioactive JA-Ile were seen to accumulate rapidly, but transiently, peaking at 1 h after the onset of salinity. This first jasmonate wave eased off completely till 24 h, when a second and higher phase of accumulation followed. Thus, the biphasic temporal pattern of *OsJAZ8* accumulation matches the biphasic temporal pattern of jasmonate accumulation.

Thus, under continued salinity, the JAZ dependent “brake” as readout of eustress *in sensu* Selye (1974) is eventually outpaced, and jasmonate signalling becomes constitutive. It is this late, deregulated phase that is maladaptive leading to distress *in sensu* Selye (1974). This distress coincides with a prolonged accumulation of ABA (Ndecky *et al*., 2023) and stomatal closure, as well as the late induction of *OsNHX1* accompanying the heavy sodium loading seen in leaf 2 (**Figs. 4, 7A**), culminating in chlorosis and necrosis of the wild type (**Fig. 1**).

Thus, the same molecule, JA-Ile, can convey two completely different signals, depending on the temporal context. Upon early and transient accumulation, JA-Ile acts as a protective signal orchestrating cellular adaptation to salinity. Upon late and constitutive accumulation, JA-Ile heralds cell death. It should be mentioned that programmed cell death, while being terminal for the individual cell can be adaptive on the level of the whole plant, because older leaves sacrifice themselves by acting as sodium sinks and, thus, shield meristem and younger leaves, such that the plant can recover, once the stress episode is over. In our study, it was mainly leaf 2 that acted as a sodium sink, thus protecting the younger leaves (**Fig. 4**), providing a further example for the adaptive role of salt-induced programmed cell death (Huh *et al*., 2002). The question, how the very same molecule, just by the difference of its temporal signature, can decide between cellular adaptation (rapid, transient jasmonate accumulation) versus cellular sacrifice for the sake of organismic adaptation (slow, but constitutive jasmonate accumulation), has been spelled out previously (Ismail *et al*., 2014). In brief, a temporal shift in early signalling events, such as calcium release and oxidative burst at the plasma membrane, can either channel to the accumulation of protective gene expression, or trigger the machinery for cellular degradation and mobilisation of resources.

This signature model predicts that engineering of a transient jasmonate signature should promote cellular adaptation and evade necrosis. This prediction has been tested experimentally by introducing a degradation-resistant dominant-negative version of OsJAZ8 under control of a salt-inducible promoter (Peethambaran *et al*., 2018). In fact, those transgenic rice plants were found to be more salt-tolerant than their respective wild type. Likewise, overexpression of CYP94C2b, a key enzyme for the catabolic degradation of JA-Ile, had the same effect (Kurotani *et al*., 2015).

How the late jasmonate accumulation mediating the cellular distress response is generated, has not been addressed, to the best of our knowledge. It is unlikely a direct response to the advent of sodium ions at the plasma membrane, but rather triggered by the intracellular damage caused by sodium ions that have penetrated through the Non-Selective Cation Channels. A prime suspect is the electron transport at the inner mitochondrial membrane, which is freely accessible for penetrated Na^+^ through the porins at the outer mitochondrial membrane and electrostatically attractive, leading to excessive superoxide production at complex III. In photosynthetically active cells, electron transport across the thylakoid is a second target. Here, the free electrons released by water-splitting at photosystem II, is a second target that is of even larger impact. Deregulated oxidative burst in these organelles might trigger salinity dependent necrosis. If so, suppression of organelle-located oxidative burst should mitigate salinity-dependent necrosis. This implication was experimentally addressed using a mitochondria-targeted Trojan peptoid carrying a coenzyme-Q type antioxidant (Asfaw *et al*., 2020). In fact, this peptoid was able to mitigate salt-induced mortality in tobacco BY-2 cells.

How this oxidative burst brings about a strong and de-regulated activation of oxylipins, remains an open question that is worth being addressed in future work. Furthermore, markers related to senescence and cell-death could be filtered out, because they should be induced during the late, but not during the early phase of jasmonate signalling.

### Conclusions and outlook

The current study explains the salt tolerance of the jasmonate-deficient *aoc* mutant by convergence of three factors, each endowed with different kinetics: a sustained stomatal conductance arising from the loss of an OPDA/JA–ABA-convergent closure signal acting on guard-cell K efflux; an early de-repression of OsHAK16-driven potassium delivery that holds the shoot K /Na balance and keeps sodium out of the youngest leaves; and, finally, the absence of the late, deregulated phase of jasmonate signalling accompanied by stomatal closure, damage-associated induction of *OsNHX1*, and chlorophyll degradation. The adaptive (“eustress”) and the maladaptive (“distress”) responses differ in their timing. The early release of *OsHAK16*, and other components of K homeostasis from jasmonate dependent suppression have to be considered as signatures of adaptation. The late induction of *OsNHX1* along with the late and constitutive activation of jasmonate signalling (also manifest by steady increase of *OsJAZ13* transcripts) are instead hallmarks of damage.

When the decision between adaptation versus necrosis depends on temporal patterns, this opens novel strategies for application, especially in the context of the accelerating secondary salinisation of major rice-growing regions of South and Southeast Asia. Rather than implementing or silencing entire signal pathways, one can try to tailor temporal patterns by introgression of promoter alleles with altered regulatory or expression features.

### Supplementary data

Fig. S1. Fresh and dry weight of rice organs in wild-type (WT) and jasmonate-deficient (*aoc*) plants under control or salt stress conditions.

Fig. S2. Potassium (K) accumulation in different organs of wild-type (WT) and *aoc* mutant plants under control or salt stress conditions.

Fig. S3. Steady-state transcript levels of ion transporter genes (*NHX1*, *HAK12*, *HAK16*, *HAK17*) in roots of wild-type and *aoc* mutant plants.

Table S1. List of primers used for gene expression (qPCR).

### Author contributions

NP: conceptualization, methodology, investigation, formal analysis, visualization, writing – original draft; MR: conceptualization, methodology, writing – review & editing; EE: resources (ICP-OES mineral analysis); PN: conceptualization, supervision, writing – review & editing.

### Funding

This work was supported by a fellowship award from the **Government of Maharashtra, India** for doctoral research at Karlsruhe Institute of Technology (KIT) to NP. Additional support was provided by a KIT–DAAD STIBET Scholarship, jointly funded by the German Academic Exchange Service (DAAD) and KIT, and by a Procope Mobility Scholarship (French Embassy / DST France) funding a bilateral Franco-German research visit to IBMP-CNRS, Université de Strasbourg.

### Data availability

All data supporting the findings of this study are available within the paper and its supplementary data published online.

## Acknowledgments

The authors thank the members of the Botanical Garden of the Joseph Kölreuter Institute for Plant Sciences (JKIP), Karlsruhe Institute of Technology, for cultivation and propagation of the rice lines. We are grateful to Antony Champion (IRD, Université de Montpellier) for providing the *aoc* mutant line, and to Thierry Heitz (Institut de Biologie Moléculaire des Plantes, CNRS, Université de Strasbourg) for critical reading of the manuscript. We thank Chantalle Kotschenreuther and Nadine Hüll (Institute of Applied Geosciences, KIT) for technical assistance with the ICP-OES analyses. We also acknowledge support by the KIT-Publication Fund of the Karlsruhe Institute of Technology.

